# Demonstrating the need for long inter-stimulus intervals when studying the post-movement beta rebound following a simple button press

**DOI:** 10.1101/2025.01.22.634263

**Authors:** Lyam M. Bailey, Timothy Bardouille

## Abstract

Voluntary movements reliably elicit event-related synchronization of oscillatory neuronal rhythms in the beta (15-30 Hz) range immediately following movement offset, as measured by magneto/electroencephalography (M/EEG). This response has been termed the post-movement beta rebound (PMBR). While early work on the PMBR advocated for long inter-stimulus intervals (ISIs)—arguing that the PMBR might persist for several seconds—these concerns have since fallen by the wayside, with many recent studies employing very short (< 5 s) ISIs. In this work we interrogated sensor-level MEG time courses in 635 individuals who participated in a cued button-pressing paradigm as part of the Cambridge Centre for Ageing and Neuroscience (Cam-CAN) project. We focussed on a subset of trials in which button presses were separated by at least 15 seconds and, using curve modelling and Bayesian inference, estimated the point at which beta power returned to baseline levels. We show that beta power takes around 4-5 seconds to return to baseline levels following movement. These results have important implications for experimental design. The PMBR is ubiquitously defined relative to a preceding baseline period; we argue that short ISIs preclude true baseline estimation and, in turn, accurate estimation of PMBR magnitude. We recommend that future studies targeting the PMBR use ISIs of at least 7 seconds—5 seconds for beta power to return to baseline, plus a 1-2 second period for proper baseline estimation. Further work is needed to clarify PMBR duration in the context of different sensorimotor paradigms and clinical populations.

---

Neuroimaging techniques such as electroencephalography (EEG) and magnetoencephalography (MEG) have long been used to study the functional properties of the human sensorimotor system noninvasively. Much of this work has focused on oscillatory neuronal rhythms in frequency ranges between 5 and 90 Hz. In particular, sensorimotor processes have long been characterized by transient changes in the Mu (8-15 Hz), beta (15-30 Hz) and gamma (30-90 Hz) frequency ranges (Cheyne and Ferrari, 2013; Pfurtscheller, 2001; Pfurtscheller et al., 1996; Pfurtscheller and Lopes da Silva, 1999; Salmelin and Hari, 1994). For example, movement or external stimulation of the hand or arm elicits suppression, or event-related desynchronization (ERD), in the mu and beta ranges (i.e., a reduction in oscillatory power relative to a preceding baseline period) (*Ibid.*). This suppression typically begins immediately prior to movement and is sustained until its offset. Approximately 500 ms following movement offset, power in the mu range returns to baseline, while beta power increases (event-related synchronization; ERS) beyond baseline levels for a short period of time—this latter response has been termed the post-movement beta rebound (PMBR). The PMBR is elicited by a range of sensorimotor processes including voluntary (Bardouille et al., 2019; Gaetz et al., 2020, 2010; Houdayer et al., 2006; Jurkiewicz et al., 2006; Labyt et al., 2003; Pfurtscheller et al., 1996) and externally induced movement (Parkkonen et al., 2015), tactile or median nerve stimulation (Bardouille et al., 2010; Gaetz and Cheyne, 2006; Houdayer et al., 2006; Parkkonen et al., 2015; Pfurtscheller et al., 2001), and imagined movement (Pfurtscheller et al., 2005; Pfurtscheller and Neuper, 1997). Multiple studies have localized the PMBR to sensorimotor cortices contralateral to movement or stimulation (Bardouille et al., 2019, 2010; Jurkiewicz et al., 2006; Pakenham et al., 2020), while a PMBR of diminished magnitude is sometimes reported on the ipsilateral side (Embury et al., 2019; Jurkiewicz et al., 2006).

Previous studies seem to agree that the PMBR following brief movements (e.g., finger extensions or button presses) or stimulation events is maximal between 500 and 1000 ms after movement (Bardouille et al., 2019; Gaetz et al., 2010, 2020; Jurkiewicz et al., 2006; Pfurtscheller, 2001; Pfurtscheller et al., 1996, 2001; Pfurtscheller and Lopes da Silva, 1999; though peak latency can increase with movement duration, Pakenham et al., 2020); accordingly, studies often target a-priori windows between 500 and 1500 ms to capture this response (e.g., Bardouille et al., 2019; Jurkiewicz et al., 2006; Pfurtscheller et al., 2001). In terms of its onset and peak, therefore, the time course of the PMBR seems well established, at least with respect to brief movements or periods of stimulation. However, the terminal point of the PMBR (i.e., when beta power returns to baseline levels) is less clear. Jurkiewicz et al. (2006) placed this point around 1 second after movement offset, while figures from other studies indicate that beta power might return to baseline levels as late as 3 or 4 seconds (see Figure 3c in Barratt et al., 2017; Figure 3 in Houdayer et al., 2006). Interestingly, some work has shown that the duration of the PMBR is affected by the amount and rate of force exerted during a voluntary movement (Fry et al., 2016), such that the PMBR may persist for as long as 7-10 seconds following offset of a sustained wrist flexion or gripping action (Fry et al., 2016; Pakenham et al., 2020). To our knowledge, no study has explicitly quantified the duration of the PMBR following a brief movement.

This issue has implications for experimental design because PMBR magnitude is ubiquitously defined relative to a baseline period (e.g., percent change, logarithm of ratio); often 1-2 seconds prior to movement onset. Depending on how closely movements / stimulations occur in time (quantified by the inter-stimulus interval; ISI) the assumption that beta power has returned to baseline from the previous trial may not hold. Thus, short ISIs may lead to improper estimation of PMBR magnitude (as discussed in Pakenham et al., 2020). Moreover, considering that multiple studies have identified PMBR magnitude as a potential biomarker for healthy ageing (e.g., Bardouille et al., 2019) as well as some clinical disorders (Gaetz et al., 2020; Xia et al., 2023; Embury et al., 2019), it is important to establish that such findings are not driven by improper baseline estimation. There is already evidence for such population differences in peak PMBR latency (Bardouille et al., 2019; Barratt et al., 2017)—differences in its duration seem likely as well, which may in turn drive apparent differences in PMBR magnitude at short ISIs.

We are not the first to consider these issues—some have proposed that an ISI of at least 9 or 10 seconds is necessary for the PMBR to return to baseline (Pakenham et al., 2020; Pfurtscheller and Andrew, 1999; Pfurtscheller and Lopes da Silva, 1999; Rhodes et al., 2024).

Nevertheless, as shown in Figure 1, many studies have used considerably shorter ISIs, often less than half (5 seconds) the aforementioned recommendations. In addition, the button-press data from the Cambridge Centre for Ageing and Neuroscience (Cam-CAN) repository (Taylor et al., 2017)—which to our knowledge represents the largest such dataset that is publicly available— includes trials with ISIs as low as 2 seconds (in fact, approximately 50% of button-press cues in this dataset occur within 3 seconds of the preceding cue; see Figure 1). All of this is to say that there is no apparent standard or minimum ISI employed when studying the PMBR.

**Figure 1.**
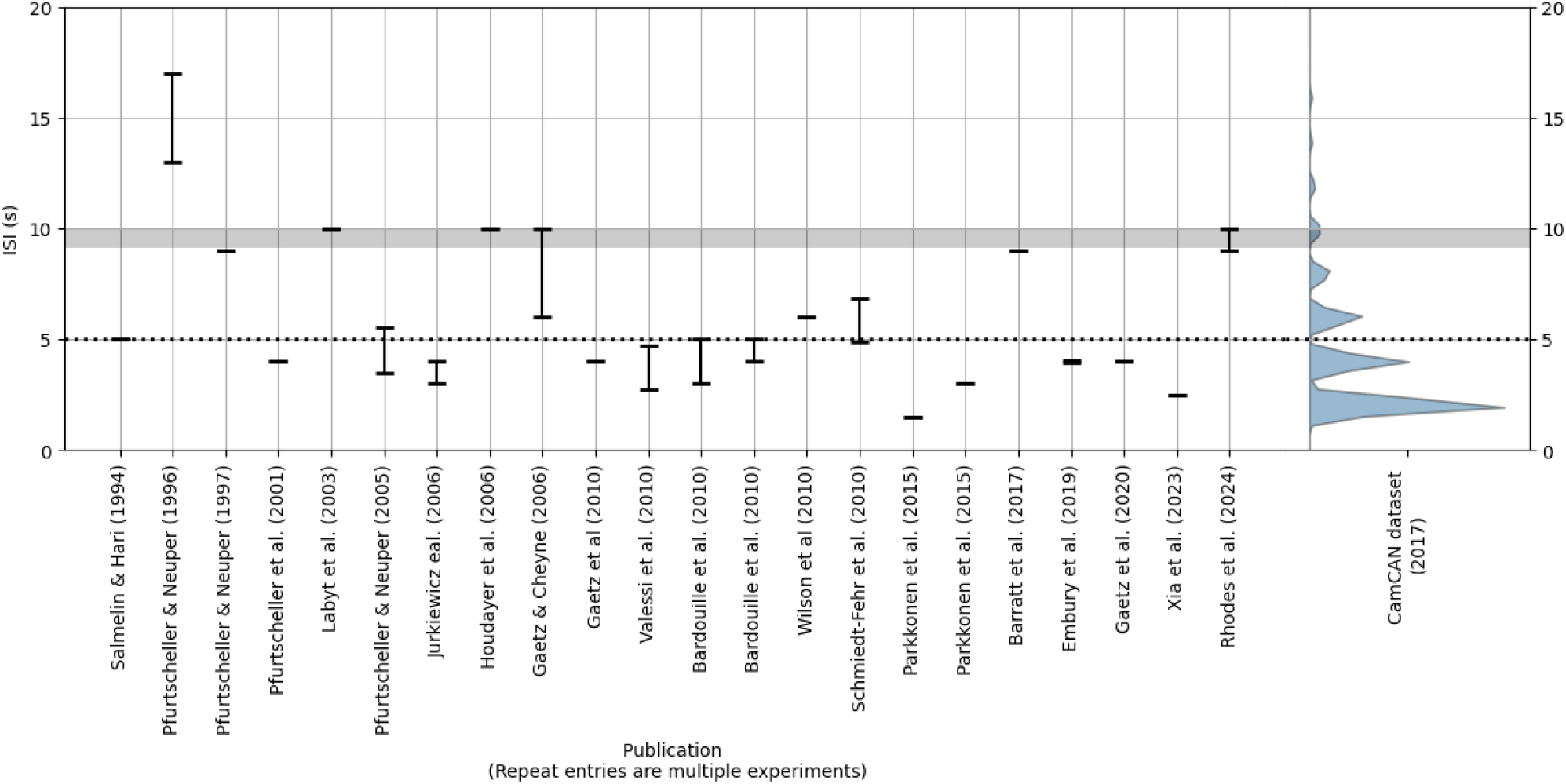
Left: Reported ISIs in a selection of the studies cited in this section. We selected studies in which participants made a single brief voluntary movement, or experienced a brief period of stimulation, on each trial. Bars represent the range of ISIs reported by the authors. **Right:** The violin plot reflects the distribution of ISIs (in this case, time elapsed since the preceding button-press cue) in the Cam-CAN button-press dataset (Taylor et al., 2017). Only three events occurred at ISIs > 20 s (not shown). The horizontal grey bar represents the 9-10 second ISI range recommended by some previous literature; the dotted line represents an ISI of 5 seconds.

The purpose of this study was to determine the minimum ISI needed to study the PMBR following a brief voluntary movement. We achieved this by approximating the time point at which beta power returns to baseline levels. We analyzed data from the Cam-CAN repository (Taylor et al., 2017), containing MEG data acquired from a large sample of participants during a simple cued button-press paradigm. We focussed on button-press events which were separated by long periods (>= 15 seconds), and considered 10 seconds post-button press to reflect a conservative benchmark for a return to baseline of the PMBR (Pfurtscheller and Andrew, 1999; Pfurtscheller and Lopes da Silva, 1999; Rhodes et al., 2024). We reasoned that PMBR duration (following a button press) would be longer than 2 seconds, given prior research practices. However, we expected that the duration would be less than the 7-10 s duration reported following wrist flexion or sustained gripping (Fry et al., 2016; Pakenham et al., 2020), given that a button-press requires less exerted force. Therefore, we predicted that the duration of the PMBR would be greater than 2 and less than 7 seconds following a button-press event. Using the Cam-CAN data clearly suggests following onto a demographic analysis of PMBR duration, however this is beyond the scope of this short communication.

## Methods

The following two sections—“Participants & Experimental Design” and “Data Collection”—have been adapted from Bardouille et al. (2019) and Power and Bardouille (2021).

### Participants & Experimental Design

A total of 708 participants (aged 18-88 years) were recruited into Phase 2 of the Cam-CAN examination of healthy cognitive ageing (Taylor et al., 2017). Of these, 650 (91.8%) had MEG data obtained during both a simple cued-button pressing task and a resting-state task.

During MEG recording each participant performed the “Sensorimotor task” (Shafto et al., 2014; Taylor et al., 2017), in which participants responded with a right index finger button press to unimodal or bimodal audio/visual stimuli. The audio stimuli were binaural pure tones of 300-ms duration at a frequency of 300, 600, or 1200 Hz. The visual stimuli were checkerboards presented to either side of a central fixation for 34-ms duration. Participants first completed a practice trial, followed by 128 trials in which 120 had bimodal stimulation, and 8 had unimodal stimulation. The order of bimodal and unimodal trials was randomized, and the inter-trial interval varied between 2 and 26 s.

We also analysed data from a “Resting state” scan, in which data were acquired for 8 min and 40 s while participants rested with their eyes closed. We confined our analysis of rest data to the middle 3-minute portion of each participant’s scan, reasoning that this period would be most representative of “true” rest.

Fifteen participants from the original sample (*N* = 650) were excluded from analyses: 10 because their raw button-press data was not available; five because Independent Component Analysis (ICA; see MEG Pre-Processing) failed to converge for their button-press data. Resting state data from one additional subject was also excluded (ICA failed to converge), but we included their button press data. Therefore, we report on 635 and 634 button-press and rest datasets respectively.

### Data Collection

Data used in the preparation of this work were obtained from the Cam-CAN repository (http://www.mrc-cbu.cam.ac.uk/datasets/Cam-CAN/) (Shafto et al., 2014; Taylor et al., 2017). MEG data were acquired at 1000 Hz with inline band-pass filtering between 0.03 and 330 Hz using a 306-channel Vectorview system (Elekta Neuromag, Helsinki, Finland). Head position was monitored continuously, and electrooculogram (EOG) and electrocardiogram (ECG) were recorded concurrently along with stimulus/response event markers.

### MEG Pre-Processing

Raw data from each subject was low-pass filtered at 40 Hz and segmented into epochs according to event selection criteria, described in “Trial Selection” below. We performed ICA on the epoched data, described previously (Bardouille et al., 2019). Finally, the MEG epochs were reconstructed from the remaining components, resulting in cleaned MEG epoch data (i.e., channels x time x epochs). We analysed cleaned epoch data from two channels, previously shown to maximally capture left (contralateral) and right (ipsilateral) beta bursts in this dataset (MEG0211 and MEG 1311 respectively).

### Trial selection

We focussed on trials with a sufficiently long period of time after each button press for the PMBR to return to baseline. To this end, we selected button press events which were followed by at least 15 seconds before the onset of the next button press. Selected events also occurred at least one second after the previous button press^1^. Including events within a few seconds of the last button press maximised the number of trials meeting our first criterion. Moreover, such events are arguably the most representative of the Cam-CAN paradigm, considering that short ISIs are very frequent (see Figure 1). Six hundred and seventeen participants had 1-4 trials meeting our criteria (611 had 1 trial), providing 626 epochs for subsequent analysis. We segmented the preprocessed data around these selected events to create “long trial” epochs of 18 seconds duration, with a 2 second pre-stimulus interval.

We also analyzed data from a comparable selection of epochs from the resting state task. Preprocessed rest data for each subject were segmented into non-overlapping 18-second epochs. We then randomly selected N epochs, where N was the number of selected button-press events for that subject.

### MEG time-frequency analysis

We computed time-frequency responses (TFRs) by applying Morlet wavelet analysis to the long trial epochs (the following was applied to the button-press and resting-state data, unless otherwise stated). Wavelet analyses were performed between 1 and 40 Hz in steps of 1 Hz, with the number of cycles equal to half the centre frequency. The data were decimated to 100 Hz to reduce processing time. This procedure returned one TFR array (time x frequency) per epoch and channel. We removed the first and last second from every TFR array to eliminate edge artifacts from the wavelet analyses. Thus, temporal structure of the final TFRs matched our trial selection criteria (i.e., 1 s before and 15 s after the button press). Finally, we averaged each TFR array over frequencies in the beta range (15-30 Hz). The result was an array of beta-band power time courses (2 channels x 626 epochs x 1600 time points). No baseline correction was applied during this process; therefore, the time courses reflected signal power (not power change) in the beta range. For the button-press data, we then calculated 1000 permuted average beta band power time courses, wherein each average comprised data from 500 randomly selected epochs. We also calculated a true grand-average (with all epochs). For the resting-state data, we calculated the average of all epochs and time points, resulting in a single point estimate of resting beta power throughout the 16 second epoch for each channel (shown in Figure 2). The results from the resting paradigm provided a reference magnitude for beta power in the resting brain, to support interpreting the grand-average button-press time courses.

**Figure 2.**
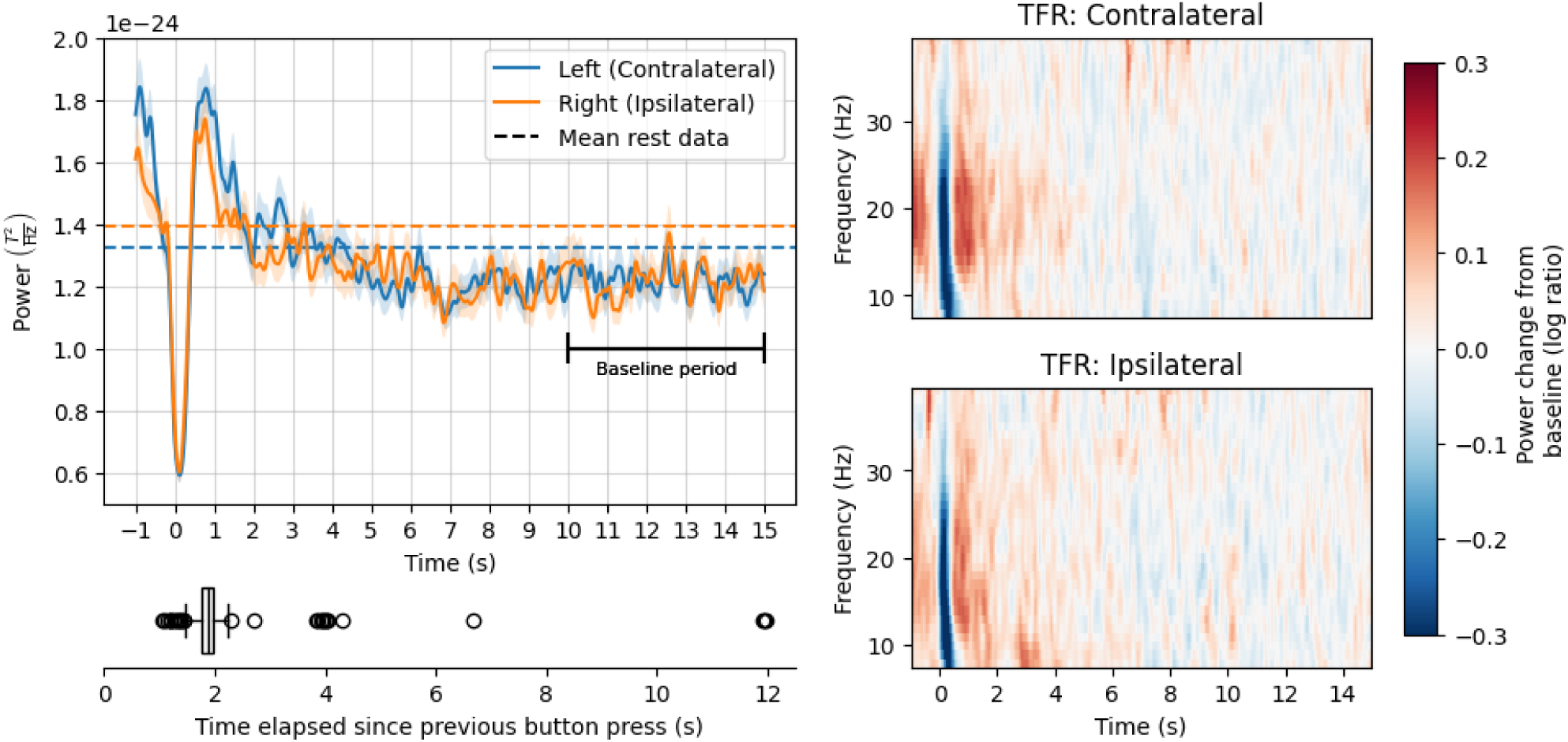
Top Left: Mean power in the beta frequency range over time, following a button press event (solid lines) or during the resting state task (dashed lines). Solid lines show grand-average time courses, averaged over epochs; dashed lines show mean power over epochs and time points. Ribbons on the solid lines are +/- 1 standard error of the mean (SE). **Bottom Left:** Distribution of time elapsed since the previous button press. **Right:** Mean power change relative to baseline (log ratio) over time and frequency, for the contralateral channel (top) and ipsilateral channel (bottom).

### Estimating the point of return-to-baseline with Weibull fitting

For each permutated beta power time course and each channel, a mean and standard deviation was calculated for the beta power in the baseline period – defined as 10-15 seconds following the button press. This time interval was selected as a baseline because it is after the longest reported PMBR duration (Pakenham et al., 2020). We then fit a modified Weibull curve (Eq. 1) (similar to previous work, e.g., Barratt et al., 2017; Liddle et al., 2016; Pakenham et al., 2020) to each time course using a least-squares minimization algorithm (implemented with the curve_fit function from the SciPy library: Virtanen et al., 2020). The Weibull curve was defined as:

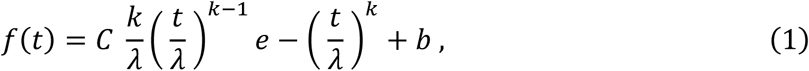

where *k* and λ were shape and scale parameters; *C* and *b* were additional parameters to further adjust to the scale and offset of our data. Curve fitting was performed on the data starting when power values surpassed the mean baseline value following suppression (i.e., PMBR onset) and ending ten seconds after the button press. Based on the fitted curve, we estimated the time at which beta power returned to baseline (*t_b_*) as the point where the fitted curve fell below one standard deviation above the mean baseline power. Over all permutations, this process returned 1000 estimates of *t_b_* per channel, with the best estimate defined as the mode *t_b_* after rounding values to two hundredths of a second.

### Estimating the point of return-to-baseline with Bayes factors

We also used Bayes factors to approximate *t_b_*. While Weibull curves have been applied in previous literature, Bayes factors provide a more formal inferential statistic. Bayes factors may provide quantitative support for observed data under a null hypothesis (Dienes, 2014; Schmalz et al., 2021; Teichmann et al., 2022) and are increasingly popular in M/EEG analyses (Grootswagers et al., 2019a, 2019b; Moerel et al., 2024, 2022; Proklova et al., 2019; Teichmann et al., 2022). A Bayes factor BF_10_ is a ratio expressing the conditional probability of some observed data under an alternative hypothesis (H_1_) over a null hypothesis (H_0_). Within this framework, BF_10_ > 1.0 and < 1.0 provides support for the data under H_1_ and H_0_ respectively, while the magnitude of BF_10_ in either direction signals the quantitative strength of evidence. We used Bayes factors to evaluate relative support for H_0_ at every time point prior to the baseline period; H_0_ predicted no difference in beta power between that time point and mean baseline power.

For each epoch, we averaged the baseline data over time to generate a distribution of average baseline power values. A Bayes factor, BF_10_, for each time point in the epoch was then computed using a paired-samples Bayes *t*-test between power at that time and the average baseline power distribution. Bayes t-tests were performed in the R (v 4.3.1) environment using functions from the BayesFactor package (Morey et al., 2022) with informed priors^2^. We estimated *t_b_* as the time point after which BF_10_ consistently favoured the null hypothesis for the remainder of the time course (i.e., BF_10_ remained below 1.0).

## Results

### Grand-average beta power time courses

Grand-average time courses for both channels, shown in Figure 2, exhibited rapid beta suppression and rebound within 1 second (s) of the button press, consistent with previous literature (see Introduction). The peak of the rebound was followed by a gradual decline in beta power before returning baseline levels. Notably, beta power in the baseline period was numerically lower than mean beta power during the resting state scan.

Beta power immediately prior to the button-press event (at -1 s) was high—around the same magnitude as the peak of the PMBR in the contralateral channel, and a little lower in the ipsilateral channel. This finding likely reflects the (gradually declining) PMBR from the preceding trial, acknowledging that most button presses in our analysis occurred within 1-2 s of the previous one (see Figure 2).

### Weibull fitting results

Results from the Weibull fitting analysis are shown in Figure 3. Over 1,000 permutations, *t_b_* estimates in both channels ranged between 2.75-6.0 s following the button press. Estimates from the two channels exhibited rather different distributions. Over 95% of estimates in the contralateral channel fell between 4.5-5.5 s, while estimates in the ipsilateral channel were more broadly distributed, with >95% between 3-5 s. The mode (i.e., best estimate of) *t_b_* for the contralateral and ipsilateral channel was 4.86 s and 4.09 s respectively.

**Figure 3.**
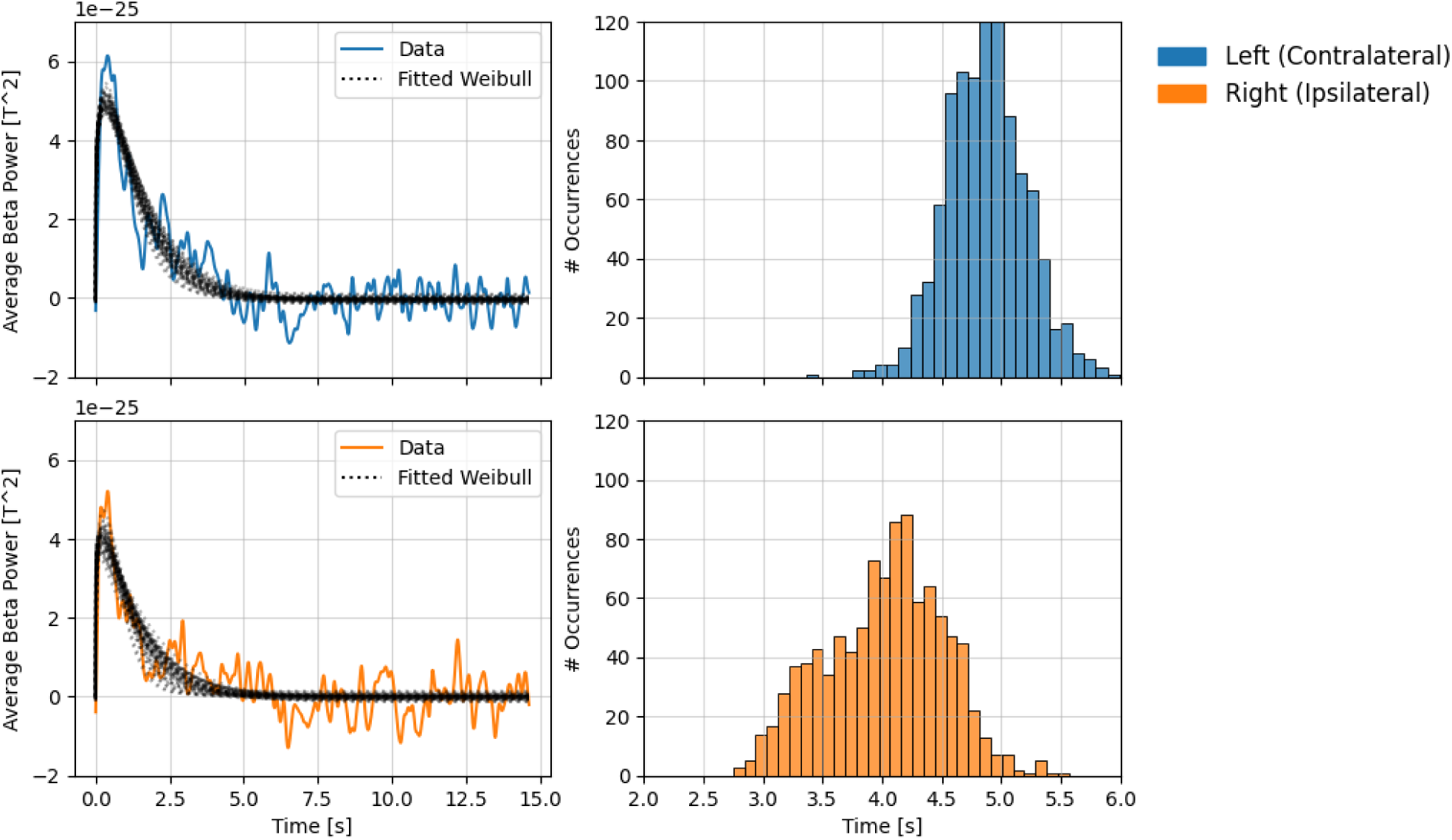
Results from the Weibull fitting analysis (1,000 permutations) for each channel. **Left:** Solid lines show grand-average beta power during the rebound period, dotted lines show permuted Weibull fits. Every 20th fit (5%) is shown here to aid visual interpretation. **Right:** Histograms show the distribution of t_b_ values obtained across permutations.

### Bayes factor results

BF_10_ time courses are shown in Figure 4. The early (and largest) peaks in each time course, around the time of the button press, correspond to the period of beta suppression. Here, beta power decreased markedly relative to the baseline period, resulting in strong evidence in favour of H_1_ in both channels. The second, smaller peak corresponds to the PMBR. Here, evidence for H_1_ was less strong (compared with the suppression period), likely because the peak of the PMBR was smaller in magnitude and exhibited larger standard errors (see Figure 2).

**Figure 4.**
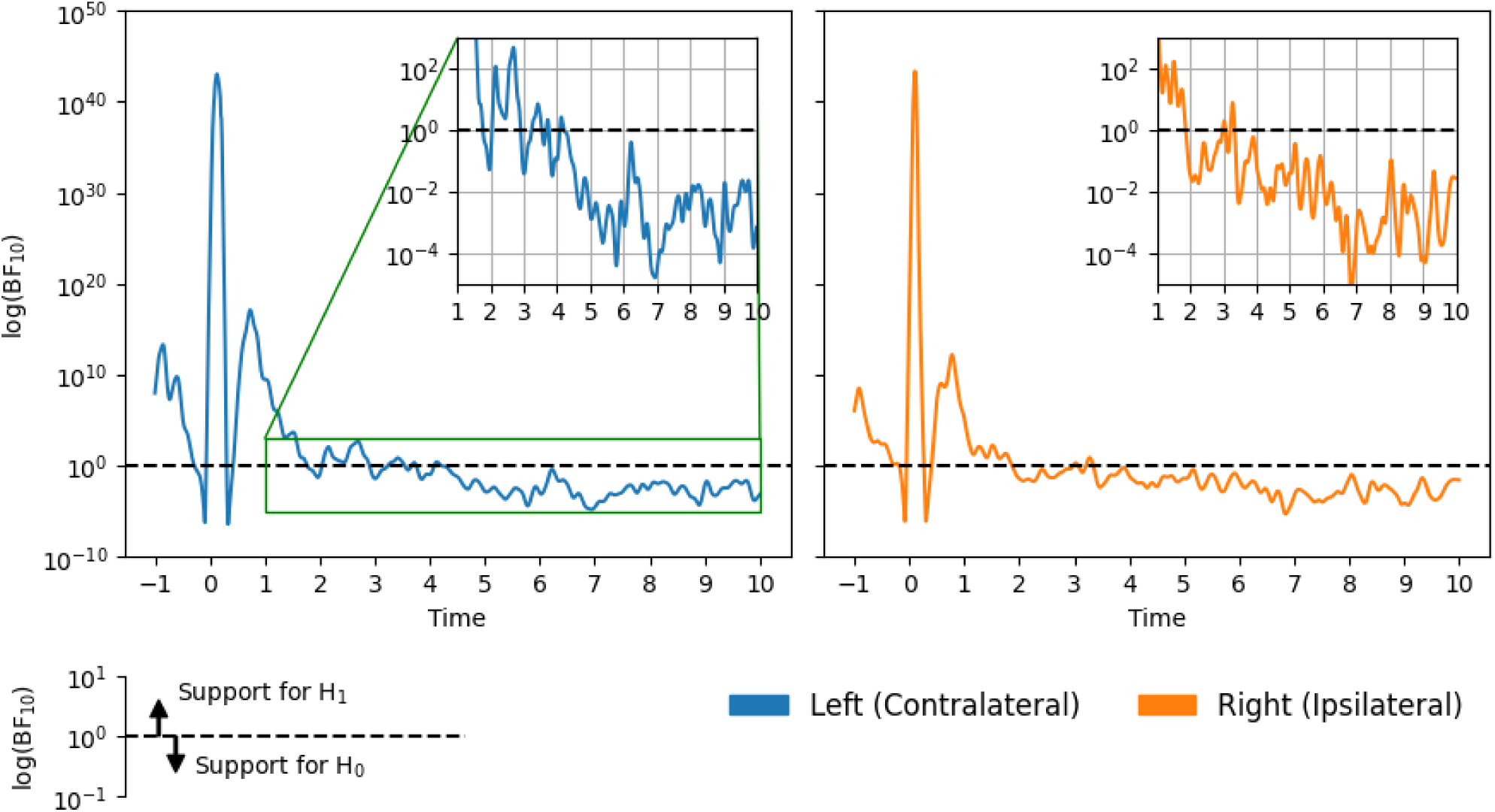
Bayes factors BF_10_ over time on a log scale, in each channel.

Of greater interest was the point at which the evidence began to consistently favour H_0_, indicating that beta power had returned to baseline. In the contralateral channel BF_10_ fell below and ceased to cross 1.0 approximately 4.25 s following the button press. Results for the ipsilateral channel indicate a slightly earlier return to baseline: around 3.25 s. These time points represent conservatively short estimates of *t*_b_ because, at these times, the quantitative support for H_0_ was relatively weak. Notably, the strength of evidence in both channels increased over time until roughly plateauing (between 10^-2^ and 10^-4^) after around 5 and 6 s on the contralateral and ipsilateral sides respectively.

Although there were small discrepancies between the Weibull and Bayes factor results (the latter provided more conservative *t*_b_ estimates), they nevertheless agree that contralateral beta power remains above baseline levels until at least 4-5 seconds following a button press. Meanwhile, ipsilateral beta power may return to baseline slightly earlier, between 3-5 seconds.

## Discussion

We investigated the duration of the PMBR following cued voluntary movement in a large sample of healthy adults. We examined a selection of trials from the Cam-CAN button-press dataset (Taylor et al., 2017) in which there was sufficient time (>= 15 seconds) following movement for the PMBR to run its course. We estimated the point *t_b_* at which beta power returned to baseline levels, defined here as average power during the 10-15 second post-movement period. Our results indicate that, following a button press, beta power on the contralateral side (where the PMBR is most reliably captured) takes at least 4-5 seconds to return to baseline levels.

These results have implications for future research on the PMBR. We recommend that researchers employ ISIs of at least 6-7 seconds: 5 seconds for beta power to return to baseline levels following movement, plus a 1-2 second period for baseline estimation. Of course, longer ISIs will provide greater certainty of accurate baseline estimation, and below we consider the possibility that certain populations may exhibit different PMBR durations, necessitating more informed experimental designs. Our recommendations are not far from (and arguably validate) those of previous work calling for ISIs of 9-10 seconds when studying sensorimotor processes with M/EEG (Pakenham et al., 2020; Pfurtscheller and Andrew, 1999; Pfurtscheller and Lopes da Silva, 1999; Rhodes et al., 2024). Moreover, we suggest that researchers analysing existing datasets (such as the Cam-CAN repository) consider excluding closely-spaced contiguous trials. An important caveat to our findings is that they only speak to the duration of PMBR following a brief voluntary movement. Considering that certain movement parameters can affect PMBR duration (Fry et al., 2016; Pakenham et al., 2020), it is unclear whether our recommendations are suitable for other sensorimotor tasks.

The need for long ISIs is particularly salient for research investigating population differences. As discussed in the Introduction, multiple studies have identified PMBR magnitude as a potential biomarker for healthy ageing and some clinical disorders (Bardouille et al., 2019; Gaetz et al., 2020; Xia et al., 2023); accurate baseline estimation is essential for such inferences. It remains to be seen whether PMBR duration differs between populations, which in turn might drive apparent differences in magnitude at short ISIs. Therefore, future studies in these areas should use sufficiently long ISIs to ensure that observed group differences in PMBR amplitude are not due to differences in the baseline. Researchers may even consider estimating the duration of the PMBR in a subset of their target population, enabling more informed experimental design.

Future research might expand on this work. For example, our findings were garnered from average responses across a wide age range (18-88 years; Taylor et al., 2017). Populations at the extremes of this range may exhibit different durations—indeed, there is already evidence for age-related changes in the temporal properties of the PMBR (Bardouille et al., 2019). Such work would benefit from examining source-localised time courses, considering that neural generators underlying the PMBR appear to shift anteriorly across the cortical surface with age (Power and Bardouille, 2021). Moreover, recent work has shown that beta ERS (which is typically measured by averaging across many trials) is mainly driven by an increase in the rate of transient beta bursts at the individual trial level (Brady et al., 2020; Little et al., 2019; Shin et al., 2017), though there is some evidence for changes to burst amplitude (i.e., the number of contributing neurons) over time (Brady et al., 2020). Future work may investigate the extent to which the protracted decrease in post-movement beta power, reported here, is driven by changes over time in burst rate and/or size of the underlying neural population. In addition, an unexpected finding from this work was that beta power during the baseline period was numerically lower than that during the resting-state task. Considering that beta ERS is broadly regarded as a marker of cortical inhibition (Pfurtscheller and Lopes da Silva, 1999), this result tentatively suggests that motor cortex is relatively disinhibited in the baseline period of an active task. Future work might investigate this possibility more thoroughly.

To summarize, we have shown that the duration of the PMBR following a brief voluntary movement is around 4-5 seconds. This estimate is considerably longer than that assumed by previous studies, many of which have used ISIs less than 5 seconds (see Figure 1), which we argue is not sufficient for proper baseline estimation. We urge caution when selecting ISIs for future experiments—our recommendation of 6-7 seconds represents, we feel, a reasonable minimum (given our data) for brief movement tasks in healthy populations. ISIs for different tasks, or studies on clinical populations, may require more judicious experimental design.

## Acknowledgements

Data collection and sharing for this project was provided by the Cambridge Centre for Ageing and Neuroscience (Cam-CAN). Cam-CAN funding was provided by the UK Biotechnology and Biological Sciences Research Council (grant number BB/H008217/1), together with support from the UK Medical Research Council and University of Cambridge, UK.

## Funding Sources

This work was supported by an NSERC Discovery grant to Timothy Bardouille (RGPIN-2018-05470)

## CRediT Author Statement

Lyam M. Bailey: Conceptualization, Data curation, Writing - original draft, Formal analysis, Investigation, Methodology, Software, Validation, Visualization. Timothy Bardouille: Conceptualization, Data curation, Funding acquisition, Project administration, Methodology, Resources, Software, Validation, Visualization, Supervision, Writing - review & editing.

## Code Availability Statement

Code for all analyses presented in this manuscript is publicly available on GitHub: https://github.com/lyambailey/PMBR_duration

## Footnotes

1 2 s was the lower end of the ISI range for this task; button presses occurring within 1 s were likely errors.

2 *t*-tests used a Cauchy prior of medium width (0.707) and a zero-point null to represent H_1_ and H_0_ respectively (default parameters in the BayesFactor package). Priors were informed in that we specified a “null interval” on the Cauchy prior—a range of non-zero values which might credibly be observed under H_0_ (Morey and Rouder, 2011; Teichmann et al., 2022). Priors without a null interval consistently yield overly conservative support for H_0_ (*ibid.*) which in this case might cause late *t*_b_ estimates. Following Teichmann et al. (2022) we determined the range of the null interval from the baseline data. We iteratively computed effects sizes (Cohen’s D) between random splits of the baseline data averaged across epochs. These effect sizes reflected changes in amplitude across time during the baseline period, which may be safely attributed to noise. Mean effect sizes following 1,000 permutations were 0.071 and 0.072 for the contralateral and ipsilateral channels respectively. These values were used as the upper limit of the null interval for each channel.

## Notes

### Competing Interest Statement

The authors have declared no competing interest.

